# AMPKi overcomes the paradoxical activation of CRAF driven by RAF inhibitors through blocking the 14-3-3 binding to its carboxyl-terminus

**DOI:** 10.1101/256651

**Authors:** Jimin Yuan, Wan Hwa Ng, Jiajun Yap, Brandon Chia, Xuchao Huang, Mei Wang, Jiancheng Hu

**Affiliations:** Division of Cellular and Molecular Research, National Cancer Centre Singapore; 11 Hospital Drive, 169610, Singapore.; Cancer and Stem Cell Program, Duke-NUS Medical School; 8 College Road, 169857, Singapore.

**Keywords:** RAF kinase, AMPK, RAF inhibitor, paradoxical activation, 14-3-3

## Abstract

The paradoxical activation of RAF kinase is the predominant challenge in cancer therapies with RAF inhibitors. The inhibitor-bound RAF molecules are able to transactivate their wild-type binding partners. 14-3-3 that binds to the carboxyl-terminus of RAF kinase has been suggested to regulate the dimer-dependent activation of RAF kinase under physiological conditions, though the molecular basis is not clear. In this study, we investigated the role of 14-3-3 in the paradoxical effect of RAF inhibitors. Firstly, we found that the 14-3-3 binding to the carboxyl-terminus of CRAF was essential for its transactivation. Further, we demonstrated that this binding enhanced the dimer affinity of CRAF. Since 14-3-3 binds to the phosphorylated motif, we next investigated and identified AMPK and CRAF itself as two putative kinases that phosphorylate redundantly the 14-3-3 binding motif of CRAF. Among RAF isoforms, CRAF plays a dominant role in the paradoxical effect of RAF inhibitors, and we thus determined whether the combinatory inhibition of AMPK and CRAF would block this effect. Indeed, our data showed that AMPKi not only blocked the RAF inhibitor-driven paradoxical activation of RAF signaling and cellular overgrowth in Ras-mutated cancer cells but also reduced the drug-resistant clones derived from BRAF(V600E)-mutated cancer cells. Finally, we showed that the 14-3-3 binding to the carboxyl-terminus of CRAF was dispensable for its catalytic function in vivo. Together, our study unraveled how 14-3-3 regulates the dimerization-driven RAF activation and identified AMPKi as a potential method to relieve the drug resistance and side effect of RAF inhibitors in cancer therapy.

## Introduction

The Ras/RAF/MEK/ERK signaling plays central roles in cell proliferation, survival, and differentiation (1-4). In normal cells, this signaling cascade is tightly regulated, and its hyperactivation causes human cancers and developmental disorders. The Ser/Thr protein kinase, RAF, is a core component of this signaling cascade, which includes three isoforms: BRAF, CRAF, and ARAF (5-7). All RAF isoforms have similar molecular structures albeit distinct traits that result in their differential ability to activate their downstream effector, MEK (8-11). Recent studies have identified RAF dimerization as a key event in the activation and regulation of this signaling cascade (12-17). In a RAF dimer, the catalysis-deficient RAF either due to mutation or inhibitor loading, facilitates its wild-type binding partner to assemble active conformation and thus triggers its catalytic activity (8). RAF dimerization not only contributes to the activation of RAF/MEK/ERK signaling under physiological and pathological conditions but also is one of important mechanisms that underlie the RAF inhibitor resistance in cancer therapy (18-20). The blockage of dimerization-driven transactivation of RAF kinase has important implications in RAF inhibitor-mediated cancer therapy.

The dimerization-driven transactivation of RAF kinase is regulated by many factors. The canonical upstream Ras activation has been shown to facilitate RAF dimerization probably through relieving the intramolecular interaction between Ras-binding domain (RBD) and kinase domain (14). Similarly, RAF inhibitors promote RAF dimerization through altering the conformation of its kinase domain (4). The dimerization-favored conformation of RAF molecule could be also achieved by catalytic spine mutations or β3-αC loop deletions that exist in cancer genomes (11,21). Furthermore, the differential molecular traits among RAF isoforms result in their distinct propensities in dimerization-driven transactivation. ARAF has a non-canonic APE motif that weakens its dimerization ability and accounts for its less activity toward MEK (11). The unique <u>N</u>-<u>t</u>erminal <u>a</u>cidic motif (NtA motif) of BRAF enables this isoform strongly transactivate the other two isoforms through dimerization, whereas the transactivation ability of CRAF and ARAF is regulated by their NtA motif phosphorylation (8). In addition, 14-3-3 has been suggested to regulate the dimerization-driven activation of RAF kinase, albeit the molecular basis of this regulation remains unknown (14,22).

14-3-3 is a family of dimeric scaffold proteins that bind to phospo-Ser/Thr within RSxpS/TxP or RxxxpS/TxP motifs (23). All RAF isoforms contain a conserved 14-3-3 binding motif in their carboxyl-terminus, though what kinase(s) is responsible for its phosphorylation remains controversial (24-27). Since CRAF is the key isoform of RAF kinase that responsible for the RAF inhibitor-induced paradoxically activation of RAF/MEK/ERK signaling in cancer therapy, in this study, we used it as a module to explore molecular mechanism that governs the 14-3-3 binding to the carboxyl-terminus of RAF kinase and thus regulates its dimerization-driven transactivation. Our data indicated that the 14-3-3 binding to the carboxyl-terminus of CRAF that is phosphorylated redundantly by AMPK and CRAF itself facilitates CRAF transactivation through enhancing its dimerization. Furthermore, we evaluated the potential value of AMPKi against the paradoxical effect of RAF inhibitors in cancer therapy and demonstrated that AMPKi could not only effectively block RAF inhibitor-induced activation of RAF/MEK/ERK signaling and cellular overgrowth in Ras-mutated cancer cells, but also reduce the formation of drug-resistant clones derived from BRAF(V600E)-mutated cancer cells. To extend our findings, we still investigated the role of 14-3-3 in the catalysis of active CRAF and found that it is dispensable for CRAF catalytic activity in vivo. Taken together, this study elucidated how 14-3-3 regulated the dimerization-driven transactivation of RAF kinase and constructed a potential approach to overcome the toxicity and side effect of RAF inhibitors in cancer therapy.

## Results

### The 14-3-3 binding to the carboxyl-terminus of CRAF regulates its dimerization-driven transactivation through elevating its dimer affinity

Previous studies have suggested that the 14-3-3 binding to the carboxyl-terminus of RAF kinase is critical for its dimer-dependent activation (14,22). To confirm this finding and further explore molecular details of this regulation, we deleted or mutated the carboxyl-terminal 14-3-3 binding motif of allosteric CRAF mutant that has a fused catalytic spine and an acidic NtA motif (DDEE/A373F) (28,29). Using these kinase-dead mutants (DDEE/A373F/S621A and DDEE/A373F/∆C), we carried out a RAF co-activation assay that developed in our previous studies (8,21). When co-expressed in 293T cell, these mutants had much less ability to transactivate the wild-type CRAF receiver and in turn induced less phosphorylation of ERK1/2 in contrast to their prototype (Figure 1A, lane 7 & 8, and 1B). This data indicated that indeed, the 14-3-3 binding to the carboxyl-terminus of CRAF plays a critical role in the dimerization-driven transactivation of CRAF.

**Figure 1.**
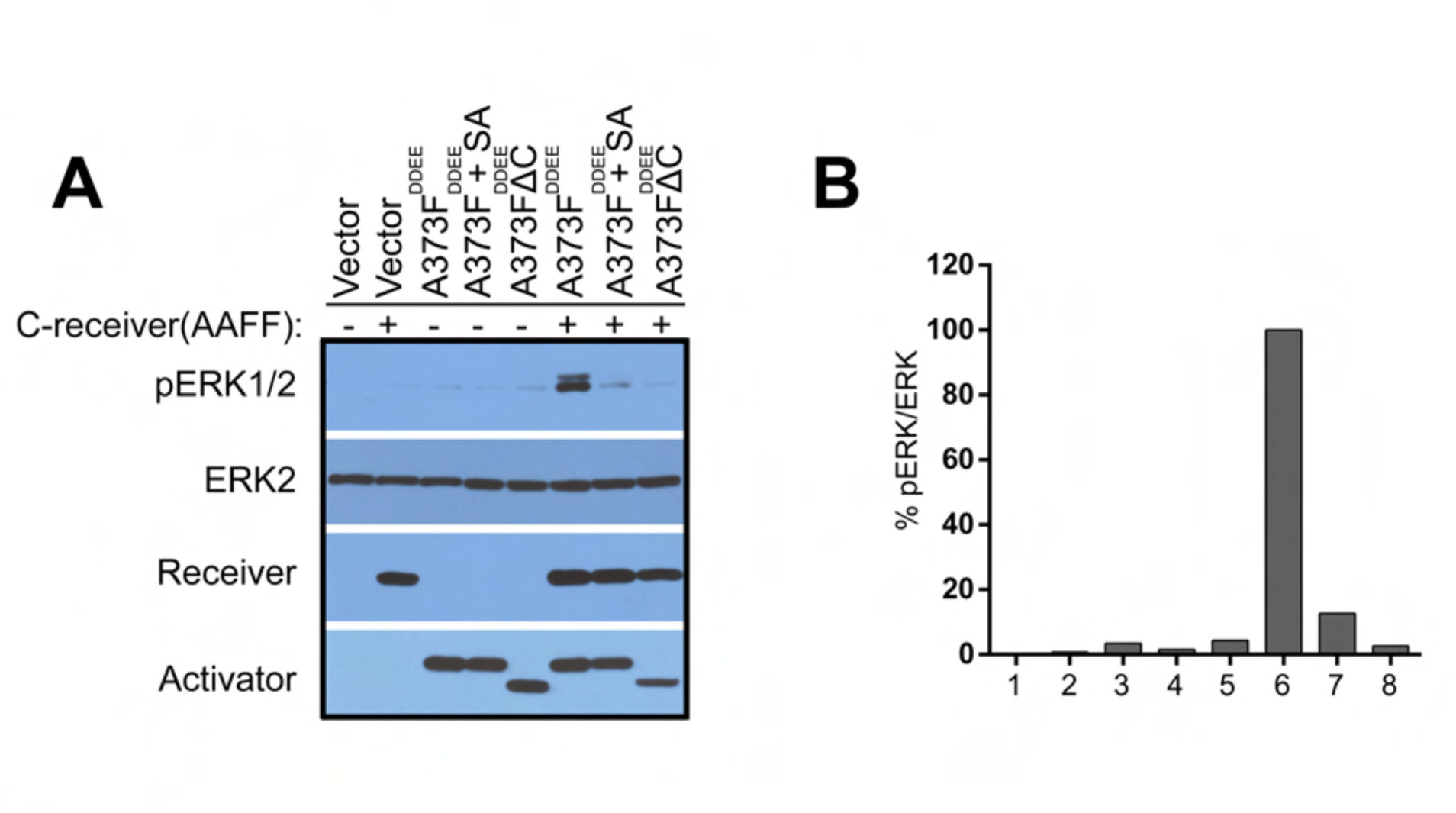
Disruption of the 14-3-3 binding to the carboxyl-terminus of CRAF blocks the dimerization-driven transactivation of CRAF. The CRAF co-activation assay was carried out as described before (8,21). Briefly, different allosteric CRAF mutants were individually co-expressed with CRAF receiver in 293T cells, and the activation of ERK1/2 was measured by anti-phospho-ERK1/2 immunoblot (A). The phospho-ERK1/2 and ERK2 were quantified by Image J and their ratio was calculated (B).

To understand how 14-3-3 regulates the dimerization-driven transactivation of CRAF, we then measured the dimer affinity of CRAF mutants with either deletion or mutation of the carboxyl-terminal 14-3-3 binding motif by complementary split luciferase assays (Figure 2A). To do this, Nluc (N-terminus of luciferase) and Cluc (C-terminus of luciferase) were fused respectively to <u>C</u>RAF <u>k</u>inase <u>d</u>omain (CKD, aa 324-648) with or without S621A mutation or deletion of 14-3-3 binding motif. Then Nluc-CKD was co-expressed with either Cluc-CKD/S621A or Cluc-CKD∆C in 293T cells (Figure 2B). The Cluc-fused R401H mutant (Cluc-CKD/R401H) that has a disrupted dimer interface was used as a control. The RAF inhibitor, vemurafenib, induced much less luciferase signals in 293T cells that express either Cluc-CKD/S621A, or Cluc-CKD∆C, or Cluc-CKD/R401H in contrast to those expressing the wild-type counterpart (Figure 2C), suggesting that CKD/S621A and CKD ∆C have impaired abilities as CKD/R401H to dimerize with CKD and the 14-3-3 binding to the carboxyl-terminus of CRAF elevates its dimer affinity.

**Figure 2.**
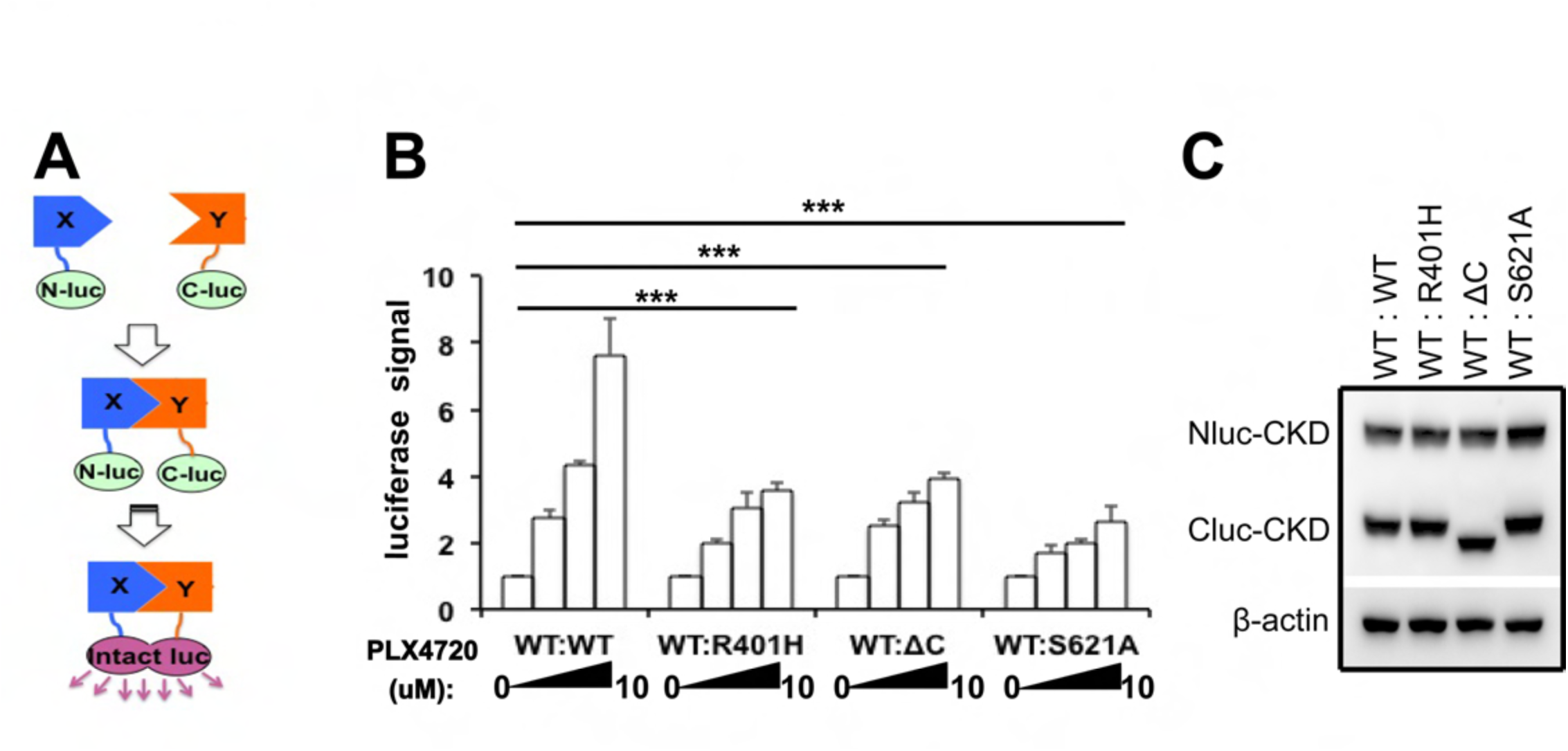
Disruption of the 14-3-3 binding to the carboxyl-terminus of CRAF inhibits RAF inhibitor-induced dimerization. The dimer affinity of CRAF or its mutants were measured by complimentary split luciferase assay as detailed in the Materials and Methods section. A, a diagram illustrating the complimentary split luciferase assay. B, Luciferase signals were measured to study the interaction between wild type and mutant CRAFs (R401H, S621A, and ∆C). Briefly, the 293T cells co-expressing Nluc-or Cluc-fused CRAF and its mutants were treated with PLX4720 before the luciferase signals were measured using illuminometer. The data is represented as the means and SD from the technical repeat of triple wells in the same study. C. The protein level of Nluc-and Cluc-CRAF fusion proteins in 293T cells from B was measured by immunoblot.

### The carboxyl-terminal 14-3-3 binding motif of CRAF is phosphorylated redundantly by AMPK and CRAF itself

It is well-known that the binding of 14-3-3 to the carboxyl-terminus of CRAF requires the phosphorylation of Ser621 in the RSxSxP motif. There exist some evidence of Ser621 phosphorylation by PKA, AMPK, or CRAF in different contexts (24-26). To clarify which kinase(s) targets this site and thus regulates the dimerization-driven transactivation of CRAF, we examined the phospho-Ser621 of wild-type and kinase-dead CRAF expressed in 293T cells with or without pharmaceutical inhibition of PKA or AMPK. As shown in Figure 3, only kinase-dead CRAF lost the phospho-Ser621 upon AMPK inhibition (lane 7), while PKA inhibition slightly enhanced the phospho-Ser621 in both wild-type and kinase-dead CRAF (lane 3 & 6), suggesting that the Ser621 in the 14-3-3 binding motif of CRAF is phosphorylated redundantly by AMPK and CRAF itself. The mechanism is unclear for the slight increase of phospho-Ser621 in the presence of PKA inhibitors. We speculate that there is a mutual inhibitory action of PKA, but further investigation is necessary for clarification.

**Figure 3.**
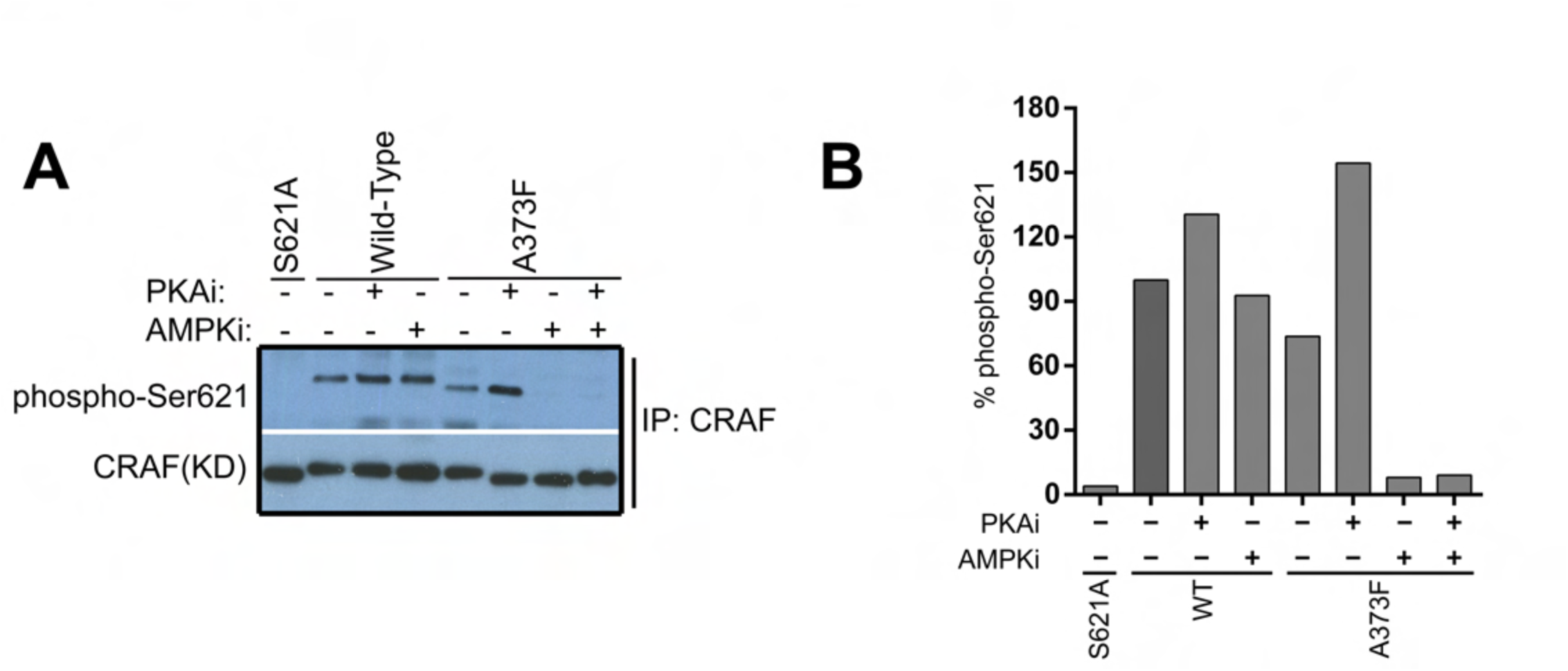
The Ser621 in the 14-3-3 binding motif of CRAF is redundantly phosphorylated by AMPK and CRAF itself, but not by PKA. Wild-type or kinase-dead mutant (A373F) of CRAF were expressed in 293T cells and treated with indicated inhibitors before purified by immunoprecipitation. The phospho-Ser621 of CRAF and its mutants was detected by immunoblot (A), and quantified using image J (B).

### AMPKi blocks the paradoxical stimulation of RAF/MEK/ERK signaling and cell growth by RAF inhibitors in Ras-mutated cancer cells

The paradoxical activation of RAF/MEK/ERK signaling driven by RAF inhibitors is not only responsible for the intrinsic resistance of Ras-mutated cancers but also one of important causes that lead to acquired resistance in BRAF(V600E)-harboring cancers (30). Moreover, CRAF has been shown as a key isoform of RAF kinase that mediates RAF inhibitor-induced paradoxical activation of this signaling pathway (18-20). As it has been demonstrated that the dimerization-driven transactivation of CRAF requires the phosphorylation of the carboxyl-terminal 14-3-3 binding motif redundantly by AMPK and CRAF itself, we next investigated whether AMPKi blocks the RAF inhibitor-induced paradoxical activation of RAF/MEK/ERK signaling in Ras-mutated cancer cells, thus represents a viable combination strategy. As reported before, RAF inhibitor, vemurafenib activated RAF/MEK/ERK signaling in a paradoxical manner in Ras-mutated cancer cell lines, H1299 (NrasQ61K) and Sk-mel-2 (NrasQ61R), but not in Ras-wlid type cancer cell line, H522 (Figure 4). This paradoxical effect of vemurafenib on H1299 and Sk-mel-2 cancer cell lines was blocked by AMPK inhibitor, Compound C (Figure 4). Furthermore, the paradoxical stimulation of cell growth by vemurafenib in H1299 and Sk-mel-2 cancer cell lines was also inhibited by Compound C (Figure 5). Noteworthy, the concentrations of Compound C used in our experiments had no apparent toxicity to these cell lines, which excludes the potential artificial arising from this AMPK inhibitor. These data further supported that AMPK-and CRAF-mediated phosphorylation of 14-3-3 binding motif plays a critical role in dimerization-driven transactivation of CRAF.

**Figure 4.**
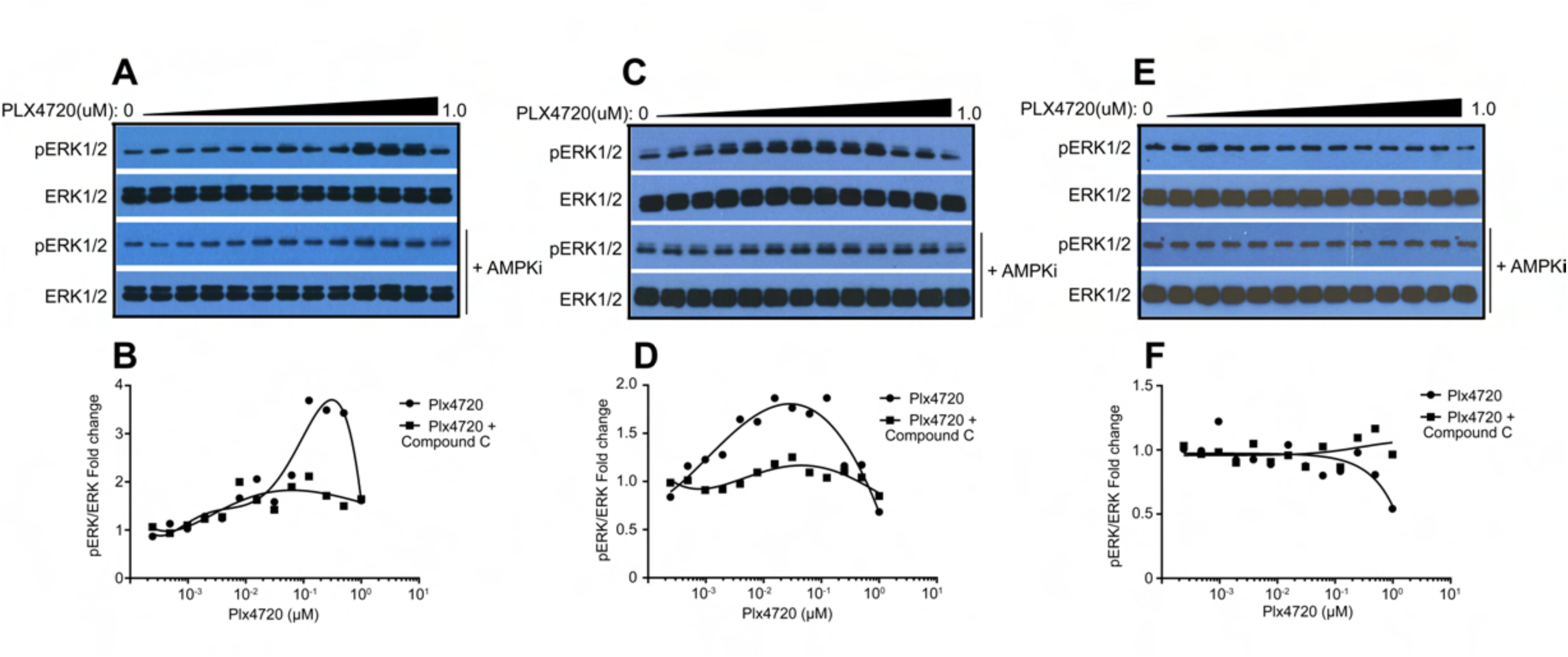
AMPK inhibitor blocks the paradoxical activation of RAF/MEK/ERK signaling induced by RAF inhibitor treatment. H1299 lung cancer cells (Nras, Q61R) (A, B), Sk-mel-2 melanoma cancer cells (Nras, Q61R) (C, D), and H522 lung cancer cells (wild type Ras) (E, F) were treated by increasing concentrations of PLX4720 with or without concurrent treatment of low doses of Compound C. The phospho-ERK1/2 and ERK1/2 were detected by immunoblots (A, C, E). The concentrations of compound C were 0.16 μM, 1.0 μM, and 0.63 μM for H1299, Sk-mel-1 and H522 cells respectively. The dose response graphs for the ratios of phospho-ERK1/2 versus total ERK1/2 measured from A, C, and E were plotted in B, D and F accordingly.

**Figure 5.**
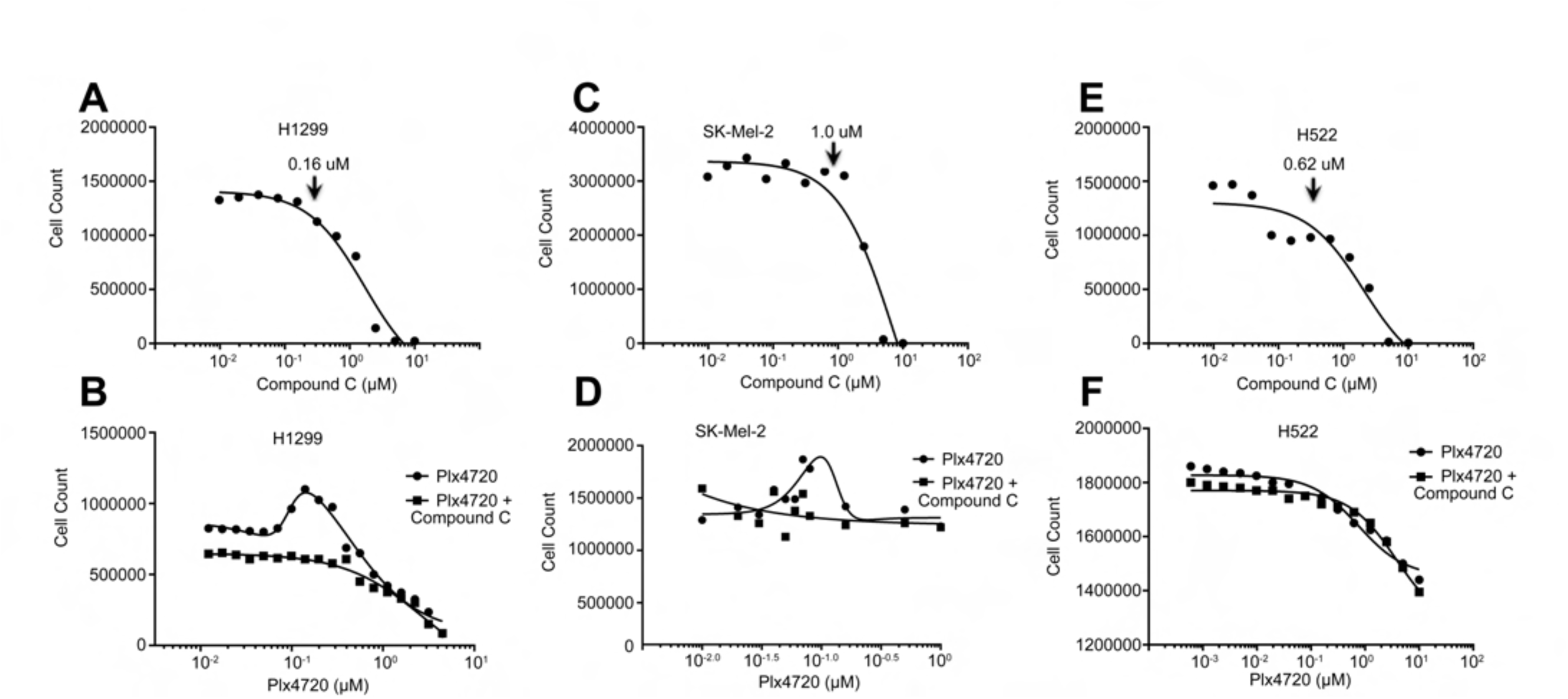
AMPKi inhibits the enhanced proliferation of Ras-mutated cancer cell lines induced by RAF inhibitor. A, C, and E, The non-toxic concentrations of Compound C for the cancer cell lines were determined by one-week culturing in a series of drug concentrations and cell counts at the end point (A, C, E). H1299 lung cancer cells, Sk-mel-2 melanoma cancer cells, and H522 lung cancer cells were tolerant to 0.16 μM, 1.0 μM, and 0.62 μM Compound C respectively. B, D, and F, The dose-dependent response of H1299, Sk-mel-2, and H522 cancer cells to PLX4720 with or without Compound C. Cancer cells were cultured in medium with 2-fold diluted PLX4720 (starting from 1.0 μM) ± 0.16 μM, 1.0 μM, or 0.62 μM Compound C for H1299, Sk-mel-2, and H522 respectively. Total cell counts were determined after one week of treatment.

### AMPKi reduces the drug resistant clones derived from BRAF(V600E)-harboring cancer cell lines

As described above, the paradoxical activation of RAF/MEK/ERK signaling also contributes significantly to acquired resistance in the treatment of BRAF(V600E)-harboring cancers with RAF inhibitors. Hence we examined whether AMPKi would enhance the efficacy of RAF inhibitors through reducing the drug resistance in BRAF(V600E)-harboring cancers. To this end, we treated A375 and A101D, two BRAF(V600E)-positive melanoma cell lines with vemurafenib alone or plus Compound C, and identified the formation of drug-resistant clones by crystal violet staining. As shown in Figure 6, the addition of Compound C at a concentration without apparent toxicity (0.62 uM) dramatically inhibited the formation of drug-resistant clones from both melanoma cell lines.

**Figure 6.**
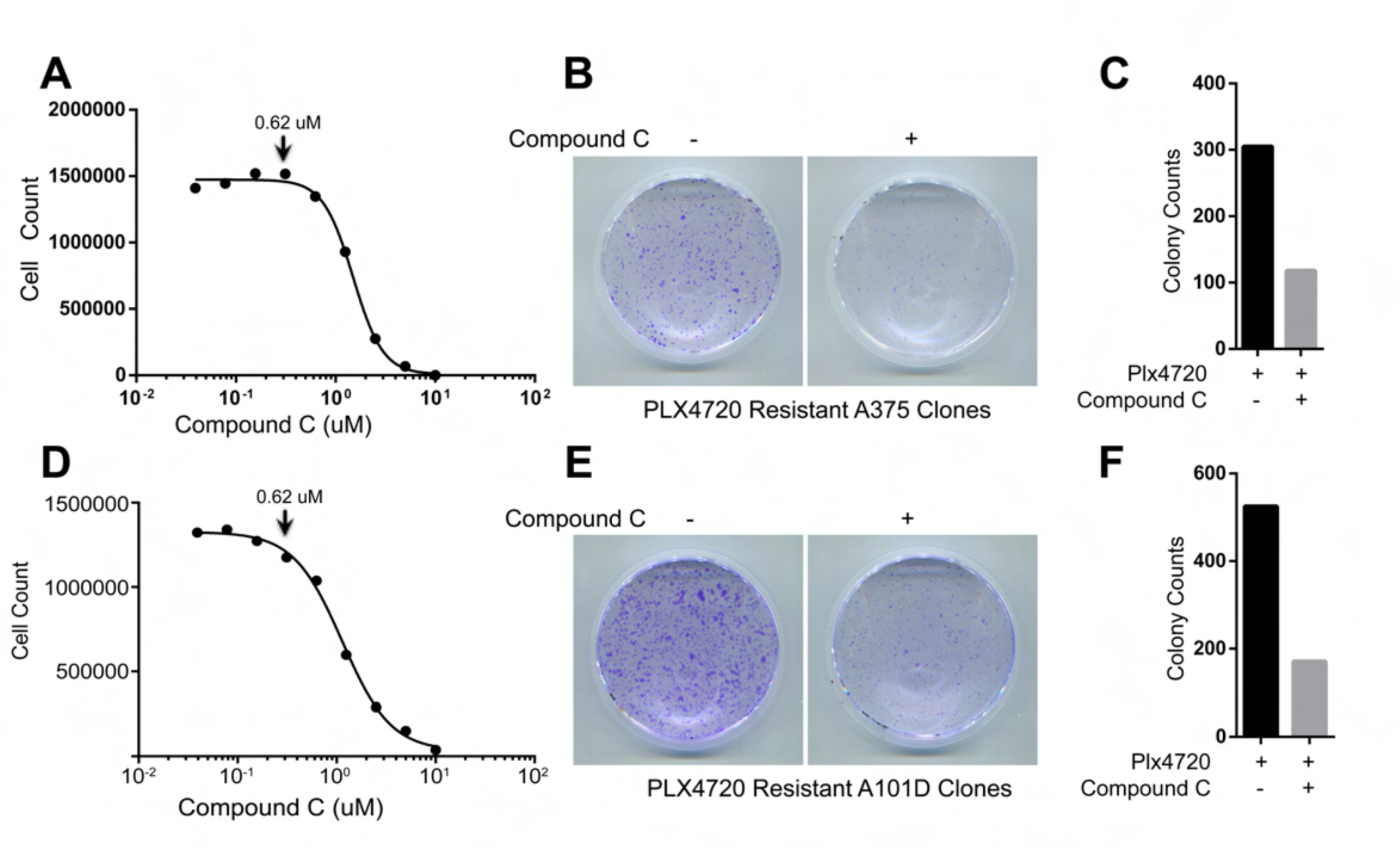
AMPKi reduces the RAF inhibitor-resistant clones derived from BRAF(V600E)-harboring cancer cells. A375 and A101D, two BRAF(V600E)-positive melanoma cell lines were treated by PLX4720 with or without 0.62 μM Compound C as detailed in the Materials and Methods section. The drug resistant clones were stained with crystal violet and counted manually. A and D, The non-toxic concentrations of Coumpound C for A375 (A) or A101D (D) melanoma cells were determined as described in Figure 5A, C, and E. Both melanoma cancer cell lines were tolerant to 0.62μM Compound C. B and E, The RAF inhibitor resistant clones derived from A375 (B) and A101D (E) melanoma cells. Cancer cells were cultured in medium with PLX4720 ± 0.62 μM Compound C as detailed in the Materials and Methods section. The PLX4720 resistant clones were identified by crystal violet staining. C and F, The quantification of PLX4720 resistant clones in B and E.

### The 14-3-3 binding to the carboxyl-terminus of constitutively active CRAF R-spine mutant elevates its dimer affinity, which is required for in vitro catalytic activity

The dimerization of RAF kinase is required for both activation and catalytic activity (11,31). Our recent study has shown that even constitutively active RAF mutants function as a dimer to phosphorylate MEK (11). In this study, we have demonstrated that the 14-3-3 binding to the carboxyl-terminus of CRAF is required for its dimerization-driven transactivation. However, whether it still regulates the catalytic activity of CRAF post activation remains unknown. To address this question, we disrupted the carboxyl-terminal 14-3-3 binding motif of constitutively active CRAF mutant (CRAF/DDEE/L397M) that generated in our previous study (21,32) and examined whether this alteration would impaire its catalytic activity. As shown in Figure 7A-B, CRAF/DDEE/L397M/S621A and CRAF/DDEE/L397M/∆C mutants exhibited a high activity as their parental protein when expressed in 293T cells. However, resembling the constitutively active CRAF/DDEE/L397M/R401H mutant with altered dimer interface (11) but not their parental protein, these mutants lost their catalytic activity in vitro upon purification, which is rescued by GST fusion (Figure 7C-D). This data suggested that CRAF/DDEE/L397M/S621A and CRAF/DDEE/L397M/∆C mutants have albeit lower dimer affinity than their parental protein though they do not associate with 14-3-3 and further supported that the 14-3-3 binding to the carboxyl-terminus of CRAF elevates its dimer affinity.

**Figure 7.**
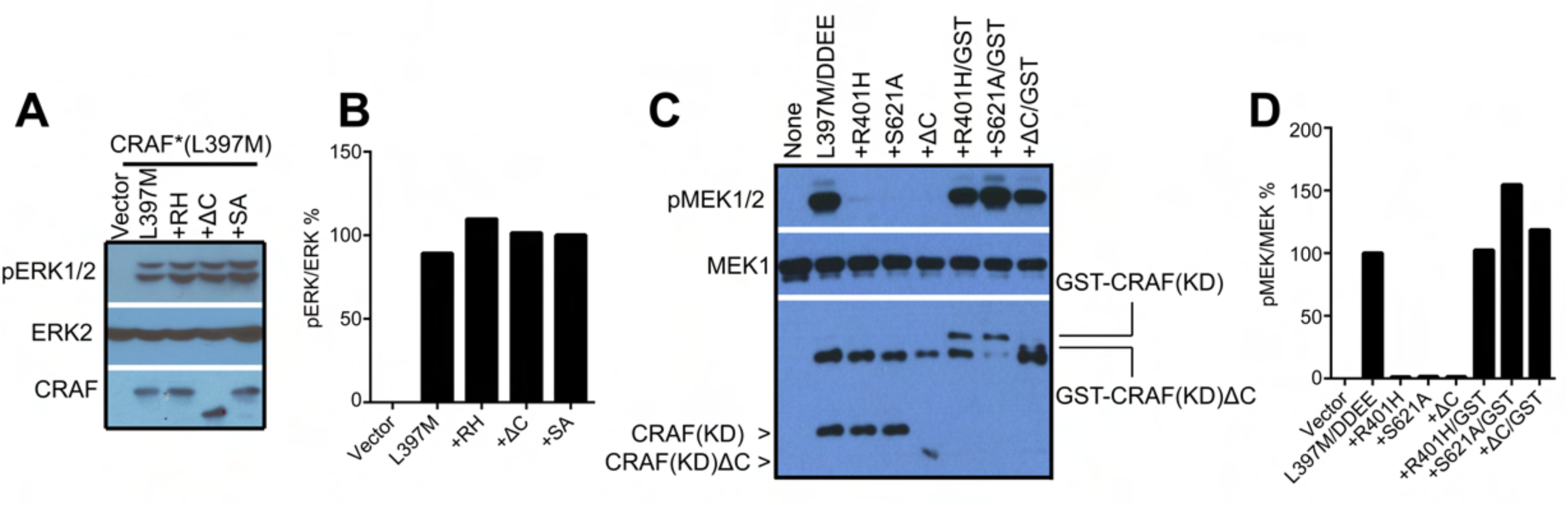
The 14-3-3 binding to the carboxyl-terminus of constitutively active CRAF mutants is not required for in vivo activity but for in vitro catalytic activity. A, The S621A mutation or the deletion of carboxyl-terminal14-3-3 binding motif does not affect the activity of CRAF R-spine mutant in vivo. CRAF mutants were expressed in 293T cells and their activity was measured by anti-phospho-ERK1/2 immunoblot. B, The ratio quantification of phospho-ERK1/2 versus ERK2 in A. C, Active CRAF R-spine mutants with S621A mutation or deletion of carboxyl-terminal 14-3-3 binding motif lose their activity in vitro upon purification, which is rescued by GST fusion. CRAF mutants in A were purified by immunoprecipitation and their activity was measured by in vitro kinase assay. D, The ratio quantification of phospho-MEK1/2 versus MEK1/2 in C.

## Discussion

The dimerization-driven transactivation plays a pivotal role in regulating the activity of RAF kinase under variable physiology and pathology conditions (33). Kinase-dead mutants or inhibitor-bound RAF molecules turn on their wild-type counterparts through this mechanism, and induce a hyperactive RAF/MEK/ERK signaling that alters cellular functions. The dimerization-driven transactivation of RAF kinase is regulated by distinct factors such as the scaffold protein 14-3-3. It has been speculated that 14-3-3 dimer binds to the carboxyl-terminus of RAF dimer and thus stabilizes RAF dimerization (14,22). This study provides strong evidence supporting for this notion. Specifically, we have shown that CRAF mutants that unable to bind with 14-3-3 have much less dimer affinity so that fail to transactivate wild-type CRAF. Furthermore, although constitutively active CRAF mutants are able to form dimers without 14-3-3 association in vivo, these dimers are so weak that dissociate during purification and thus lose catalytic activity in vitro. These evidence are consistent with our recent findings from oncogenic RAF mutants with β3-αC loop deletions, which suggest that a strong dimerization drives transactivation of RAF kinase whereas a weak dimerization is required for catalytic activity following its activation (11).

14-3-3 binds to the RSxSxP or RxxxS/TxP motif only when its Ser/Thr is phosphorylated (23). However, the protein kinase(s) that phosphorylates the 14-3-3 binding motif in the carboxyl-terminus of RAF kinase was not clearly defined prior to this study, despite the potential involvement of PKA, AMPK, or RAF (24-27). Using kinase-dead mutants and pharmaceutical inhibitors, we have demonstrated that the carboxyl-terminal 14-3-3 binding motif of CRAF isoform is redundantly phosphorylated by AMPK and CRAF itself but not PKA, in cell lines we studied. This conclusion is further supported by the finding that AMPK inhibitor blocks the paradoxical activation of RAF/MEK/ERK signaling driven by RAF inhibitor in cancer cell lines with active Ras mutations.

The paradoxical activation of RAF/MEK/ERK signaling leads to both the intrinsic resistance in Ras-mutated cancers and the acquired resistance in BRAF(V600E)-driven cancers, which severely limits the efficacy of RAF inhibitors in cancer therapy (30). Moreover, it also induces the secondary tumors in RAF inhibitor-treated patients. Our finding that AMPK inhibitor abolishes the RAF inhibitor-driven paradoxical activation of RAF/MEK/ERK signaling in Ras-mutated cancer cells and thus inhibits their overgrowth provides a potential combination therapy to control Ras mutation-driven cancers. In addition, since the combination of AMPK inhibitor with RAF inhibitor also reduces dramatically the drug-resistant clones derived from BRAF(V600E)-harboring cancer cells, it will improve the treatment of this type of cancers with RAF inhibitors.

## Materials and Methods

### Chemicals and Antibodies

Antibodies used in this study included: anti-phosphoERK1/2 (#4370), anti-phosphoMEK1/2 (#9154), and anti-MEK1/2 (#9124) (Cell Signaling Technology); anti-phosphoSer621 (AM00131PU-N, Acris antibodies)(34); anti-FLAG (F1804, Sigma); anti-HA (#4810, Novus Biologicals); anti-ERK2 (SC-154, Santa Cruz Biotechnology); anti-ERK1/2 (A0229, AB clonal); and HRP-labeled secondary antibodies (#31460 & #31430, Invitrogen). PLX4720 (A-1131) was purchased from Active Biochem; H-89 (S1582) and Compound C (S7306) from Selleckchem; and D-luciferin (LUCK-2KG) from Gold Biotechnology. All other chemicals were obtained from Sigma.

### Plasmids, Cell lines and Protein Expression

All expression vetcors used in this study were either purchased from Addgene or synthesized by Integrated DNA Technologies, as noted in the text. All mutatants were generated by PCR, tagged with either FLAG or HA or His, and cloned into vectors by Gibson assembly. pCDNA3.1(+) expression vector (Invitrogen) was used for transient expression in mammalian cells and pET-28a (Novagen) for bacterial expression.

Cancer cell lines: H1299, Sk-mel-2, H522, A375, and A101D were obtained from ATCC. All cancer cell lines were maintained in DMEM medium with 10% FBS (Hyclone). For exogenous expression, 293T cells were transfected with the appropriate plasmids using the Biotool transfection reagent by following the manufacturer’s protocol.

6xhis-tagged MEK1 (K97A) was expressed in bacteria BL21(DE3) strains and purified by using Nickel column (Qiagen) and following our previous protocol (35).

### Complementary Split Luciferase Assay

293T transfectants that express different pairs of Nluc-and Cluc-fused CRAF or CRAF mutants were plated in 24-well Krystal black image plates at the the seeding density of 2x10^5^ per well. 24 hour later, D-luciferin (0.2mg/ml) and PLX4720 (0, 2.5, 5,10μM) were added to the culture; the incubation was allowed for 30 min before the luciferase signals were measured by Promega GloMax^®^-Multi Detection System.

### Immunoprecipitation, *In Vitro* Kinase Assay, and Western Blotting

Immunoprecipitations were performed as described previously (8). Briefly, whole-cell lysates were mixed with either anti-HA, or anti-FLAG beads (Sigma), rotated in cold room for 60 min, and washed three times with RIPA buffer. For *in vitro* kinase assays, the immunoprecipitants were washed once with kinase reaction buffer (25 mM HEPES, 10 mM MgCl2, 0.5 mM Na3VO4, 0.5 mM DTT, pH 7.4), then incubated with 20μl kinase reaction mixture (2 ug substrate and 100 mM ATP in 20μl kinase reaction buffer) per sample at room temperature for 10 min. Kinase reaction was stopped by adding 5μl per sample 5XLaemmli sample buffer. Immunoblotting was carried out as described before (36). All blots were quantified by using Image J and the graphs were generated by using GraphPad Prism 6.

### Drug Toxicity Test and Cell Proliferation Assay

Cells were seeded in 6-well plates by 2x10^5^ per well one day before treatment. Compound C was added to cell cultures as contiguous concentrations from 10 μM with 2-fold dilutions. One week later, cells were harvested and counted by using hemocytometer. The highest concentrations of Compound C that have no apparent toxicity on cell growth were applied to cell proliferation assays or clone formation assays below.

To examine the effect of PLX4720 on cellular proliferation, cells were seeded by 5x10^4^ per well in 6-well plates and PLX4720 was added to cell cultures as contiguous concentrations from 1 μM with 2-fold dilutions together with or without constant concentrations of Compound C according to the results from drug toxicity tests. The culture medium with drugs was changed every other day, and 6 day later, cells were harvested and counted as above. The cell growth curves were generated by using GraphPad Prism 6.

### Clone Formation Assay

Cells were seeded in 60-mm dishes at the density of 2x10^5^ per dish. 24 hours later, PLX4720 (0.2μM) was added cell cultures together with or without Compound C (0.62μM). The culture medium was changed every other day and the concentration of PLX4720 was increased by 2-fold every time until it reached 0.8μM. After two weeks culturing, cells were fixed with 4% paraformaldehyde and stained with crystal violet solution. The numbers of clones were counted manually and the graphs were generated using GraphPad Prism 6.

## Author Contributions

J.Y. and J.H. designed the study; J.Y., W.H.N, J.J.Y., B.C, X.H. and J.H. carried out experiments; M.W. and J.H. supervised all experiments and interpreted experimental data; J.H. wrote the manuscript; M.W. revised manuscript; and all authors commented and approved the manuscript.

## Acknowledgement

We thank the laboratories of Dr. Sabapathy and Dr. Virshup for their help in experimental technologies. This study is supported by NCCRF startup grant (NCCRF-SUG-JH), NCCRF bridging grant (NCCRF-YR2016-JUL-BG1), NMRC seeding grants (NCCSPG-YR2015-JUL-14 and NCCSPG-YR2016-JAN-17), Duke-NUS Khoo Bridge Funding Award (Duke-NUS-KBrFA/2017/0003), SingHealth Fundation startup grant (SHF/FG692S/2016), and Asia Fund Cancer Research (AFCR2017/2019-JH). The authors declare no potential conflicts of interest.

